# Pose-derived microstructure resolves how hunger and social isolation shape social–food choice in male mice

**DOI:** 10.64898/2026.11.12.765913

**Authors:** Aryan Atul Surana, Niranjan Biju, Soumyajyoti Pal, Inga D Neumann, Virginie Rappeneau

## Abstract

Preference indices reduce choice between competing rewards to a single score, obscuring how internal state and experience shape the behaviour that produces it. We combined the social versus food preference test (SFPT) with markerless pose estimation, cross-validated decoding using markerless pose estimation, and Zif268 immunofluorescence in adult male CD-1 mice that were group-housed (GH) or single-housed (SH) and tested fed ad libitum (AL) or after 16 h of food deprivation (FD). Food deprivation shifted preference toward food in both housing conditions, whereas SH shifted it toward the social stimulus. In GH mice, the food shift reflected longer food-investigation bouts and stronger food-directed orientation, but not more food-chamber entries. SH-FD mice showed no significant preference, yet they investigated both stimuli and approached them at high speed. Pose-derived features decoded feeding state (AUC 0.94) and housing condition (AUC 0.85), whereas manually scored measures decoded feeding state (AUC 0.84) but not housing (AUC 0.56). In exploratory analyses, simultaneous choice between food and social stimuli was associated with higher Zif268 expression in the basolateral and central amygdala and supraoptic oxytocin neurons of GH-FD mice, and in the BNST and hypothalamic oxytocin neurons of SH-AL mice. Pose-derived microstructure thus resolves components of social–food choice that a single preference score merges.

**Highlights:**

- Null preference scores conceal distinct social–food strategies.
- Food deprivation prolongs food visits, not chamber entries.
- Single-housed, food-deprived mice show no preference but approach both stimuli fastest
- Pose features decode housing history; manual scores do not.

## 1. Introduction

Animals continually allocate limited time and energy among competing needs, and the outcome depends on internal state, recent experience and context as much as on the value of either option [1,2]. Hunger redirects behaviour toward food at the expense of other drives [3–6], and social isolation induces a distinct state that alters subsequent social and affective behaviour [7,8]. In humans, acute isolation evokes midbrain responses to social cues that resemble those evoked by food cues after fasting [9]. Yet, most behavioural assays summarise such choices as a single preference score. A score reports which option prevailed, but not how: through frequent brief visits or a few sustained bouts, through rapid approach or prolonged orientation without approach. Identical scores can therefore conceal different strategies, and a null score can conceal active engagement with both options rather than indifference.

The social versus food preference test (SFPT) is a tractable assay for this problem, because social interaction and feeding are both salient, and their prioritisation is sensitive to physiological and environmental conditions [10–12]. Food deprivation (FD) shifts preference from social interaction toward food in C57BL/6J mice [10,12,13] and, with longer FD, in rats [10,12]. Housing also matters: after 24 h of FD, group-housed (GH) female rats preferred food, whereas single-housed (SH) females preferred social interaction [14]. However, these studies rely largely on investigation times and preference indices, and recent work shows that motivated behaviour has a temporal structure that such indices compress [10,14,15]. In mice facing conflicting hunger and thirst, choices were organised into persistent bouts with stochastic switches, modelled as a goal state diffusing across an energy landscape in which each need defines a “well” that deepens with the strength of the need and flattens as it is met [15]. A preference thus reflects not only which well an animal enters, but how deep it is and how long the animal stays in it, which a single index cannot separate. It therefore remains unclear whether feeding state and housing history influence behavioural prioritisation through shared or distinct mechanisms.

Markerless pose estimation and the use of DeepLabCut technology [16–18] now enables continuous, high-resolution quantification of posture and movement, capturing how animals approach, orient towards and engage with a stimulus rather than simply how long they remain nearby. This offers an opportunity to distinguish the behavioural effects of feeding state and housing history that may be obscured by a single preference score. It may also reveal changes in behavioural strategy that conventional scoring fails to detect, even when overall preference remains unchanged. To our knowledge, pose-derived features have not previously been used to decode internal states during the social versus food preference test.

However, the neuronal circuits engaged in the choice between social interaction and food remain poorly understood. Several brain regions implicated in social and feeding behaviours may contribute to this choice [19,20]. Hypothalamic hunger circuits bias behaviour toward food at the expense of competing drives [3–6,13]. In the amygdala, the basolateral nucleus (BLA) contains largely nonoverlapping representations of social and sucrose reward [21] and genetically defined neurons that control both food consumption and social interaction [22]. The central amygdala (CeA) promotes food intake through a positive-valence mechanism [23]. Other nodes carry overlapping or interacting representations of the two reward types, including orbitofrontal ensembles [24] and ventral tegmental dopamine neurons [25]. Two further systems are particularly relevant to a choice that engages both hunger and isolation. The bed nucleus of the stria terminalis (BNST), an extended-amygdala structure, regulates anxiety-related and motivational states [26] and may be recruited when competing needs conflict. Oxytocin neurons of the paraventricular (PVN) and supraoptic (SON) nuclei are activated by social stimuli, and oxytocin promotes social behaviour [27], restrains feeding [28], and has been recently implicated in the isolation-dependent reorganisation of social–food priorities in rats [14], which makes the neuropeptide a likely candidate integrating both needs. Whether these regions are recruited by the choice itself, rather than by either stimulus alone, remains unclear. Zif268/EGR1 (early growth response 1) immunoreactivity provides a cellular activity marker [29,30], and comparing its expression after dual-stimulus choice with that after each stimulus alone allows to identify regions engaged by both stimuli.

Here we tested adult male CD-1 mice in the SFPT in a 2 × 2 design of housing (GH or SH) and feeding state (ad libitum, AL, or 16 h of FD), and compared conventional behavioural scoring with markerless pose estimation. We hypothesised that (1) hunger and social isolation influence social-food prioritisation in different ways; (2) pose estimation captures more about feeding state and housing history than manual scoring; and (3) choosing between social and food stimuli engages specific brain regions depending on the animal’s state.

## 2. Materials and Methods

### 2.1. Animals

Male CD-1 mice were obtained from Charles River Laboratories (Sulzfeld, Germany) at 8 weeks of age. On arrival, mice were housed in groups of three to four in standard cages measuring 42 × 22 × 14 cm under controlled laboratory conditions (22 °C, 60% humidity, 12-h light/dark cycle; lights on at 07:00). Standard chow and water were available ad libitum unless stated otherwise. Age- and sex-matched CD-1 males served as social stimuli. Procedures followed ARRIVE 2.0 guidance [31], and international regulations for the care and use of animals (Directive 2010/63/EU), with ethical approval from the government of Unterfranken (RUF-55.2.2-2532-2-2082).

### 2.2. Experimental design and behavioural tests

After two weeks of acclimation and daily handling, mice were assigned to group housing (GH; three to four per cage) or single housing (SH; 26 × 20 × 14 cm cages) for two weeks before testing. Mice were habituated to Honey Pops (Kellogg’s) in the home cage for 2–3 days and then underwent eight 5-min habituation sessions in the apparatus across three days (reward side counterbalanced) (see Supplementary Methods S1). Mice were then tested either fed ad libitum (AL) or food-deprived (FD) for 16 h overnight. Body weight was recorded before and after deprivation. The groups were GH-AL (n=10), GH-FD (n=10), SH-AL (n=11) and SH-FD (n=18). The SH-FD group was larger to account for its greater behavioural variability. The main cohort underwent the SFPT and immunofluorescence; separate cohorts underwent the social-only (ST) or food-only (FT) tests.

All behavioural tests, i.e., the SFPT, ST and FT tests were performed between 08:00 and 13:00 under dim light (≤15 lux) in a three-chamber apparatus (82 × 82 × 41 cm) comprising two end chambers connected by a neutral central chamber through openings (3.5 × 4 cm). One wire-mesh stimulus cage (7 × 7 × 6 cm) was placed in each of the end chambers. In the SFPT (adapted from [10–12]; Fig. 1B), the experimental mouse first explored two empty cages located in the end chambers for 5 min. Thereafter, the 2 empty cages were replaced by one cage containing an unfamiliar male conspecific and another one containing Honey Pops (identity and side counterbalanced), for 10 min. Stimulus-directed investigation was defined as the nose entering a predefined interaction zone. The preference index was calculated as (social − food)/(social + food); positive values indicate a social bias whereas negative values indicate a food bias. Behaviour was scored automatically with DeepLabCut [17,33], from which behavioural features were computed using custom Python scripts (Section 2.3). For comparison, videos were also scored manually by observers blinded to housing and feeding state using the Behavioral Observation Research Interactive Software (BORIS) [32].

**Figure 1.**
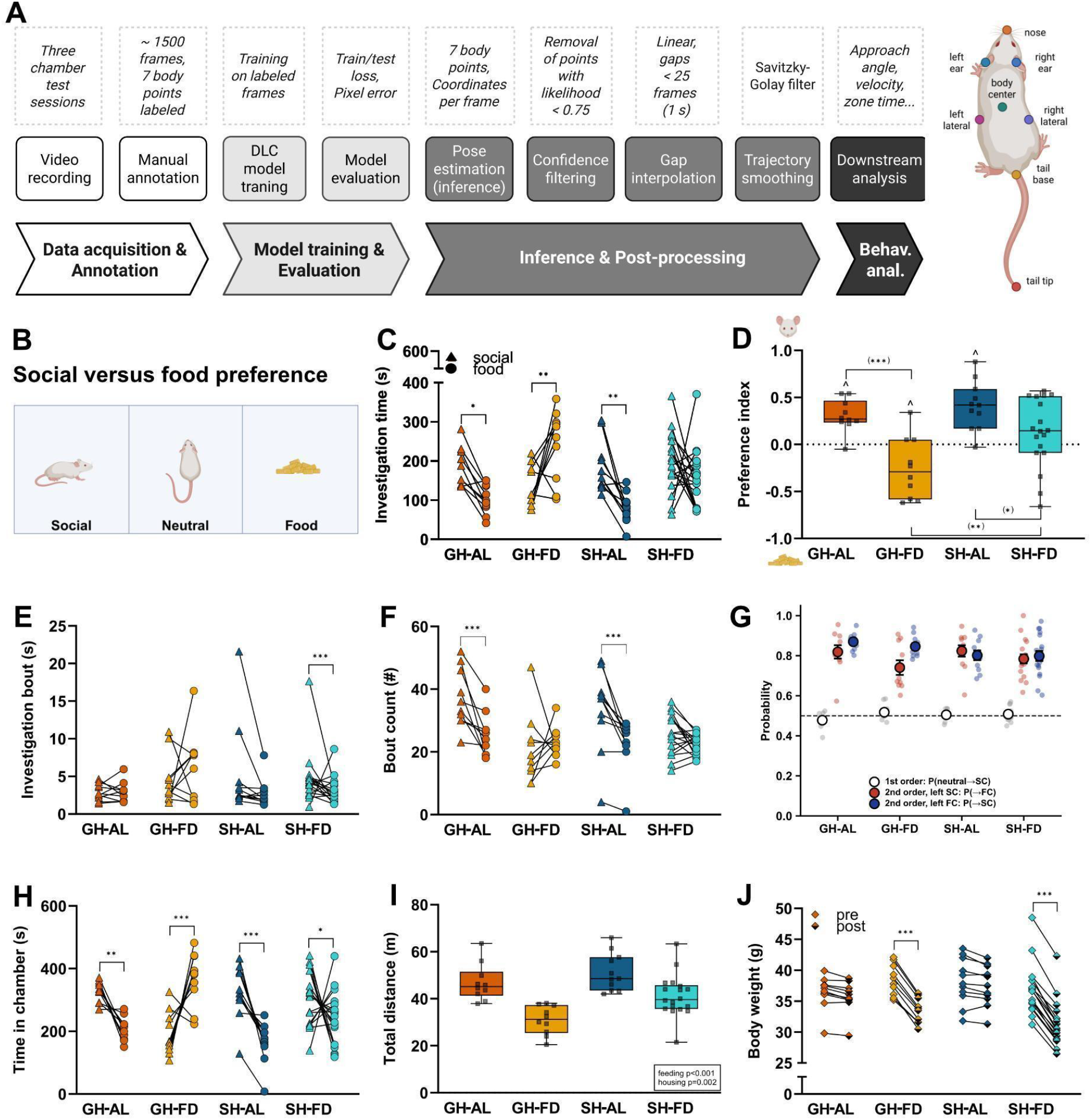
Feeding and housing conditions bias the choice between social and food rewards. (A) DeepLabCut workflow. (B) Social versus food preference test design. (C) Investigation time toward the social and food stimuli in GH-AL, GH-FD, SH-AL and SH-FD mice. (D) Preference index (positive: social preference; negative: food preference). (E) Duration and (F) number of investigation bouts. (G) First and second order transition probabilities for chamber transitions. (H) Time in the social and food chambers. (I) Total distance. (J) Change in body weight before and after the social versus food preference test. Data are presented as individual data points or box-and-whisker plots (minimum to maximum, all points). Statistical analysis: *p<0.05, **p<0.01, ***p<0.001 (C), (E), (F), (H) and (J) (Two-way repeated-measures ANOVA, Tukey post hoc); (D) and (I) (Two-way ANOVA, Tukey post hoc); (*)p<0.05, (*)*p<0.01 (D) (planned separate comparisons using Student’s t-test); ^p<0.05 (D) (one-sample t-tests versus 0). Sample sizes: GH-AL, n=10; GH-FD, n=10; SH-AL, n=11; SH-FD, n=18. Abbreviations: AL, fed ad libitum; FD, food-deprived; GH, group-housed; SH, single-housed. (A,B) Created in BioRender.

In the ST, an unfamiliar conspecific was placed in a stimulus cage, and the other cage remained empty. In the food-only test (FT), Honey Pops were placed in one cage and the other remained empty. Reward value was further assessed using a conditioned place preference (CPP) paradigm, in which Honey Pops and standard food were paired with distinct chambers (see Supplementary Methods S1). Videos from ST, FT and CPP sessions were only scored manually using BORIS [32]. As these experiments were intended solely to confirm the rewarding properties of the stimuli, analyses were limited to investigation time and frequency (or time spent in each chamber for CPP).

### 2.3. Pose estimation and feature extraction

Behaviour was quantified from videos of the SFPT (Section 2.2) Videos were analysed with DeepLabCut v3.0.0rc9 [17,33], tracking eight key points (nose, ears, body centre, lateral body points, tail base and tip; Fig. 1A). The network was trained on 2,096 frames from 78 SFPT videos (1,985 training, 111 test). Accuracy was evaluated using COCO Object Keypoint Similarity (OKS), which measures keypoint precision relative to animal body size and body-part variability. Averaged across standard thresholds (0.50–0.95), the model achieved a mean Average Precision (mAP) of 95.8% and a mean Average Recall (mAR) of 96.7% on held-out frames. Estimates with likelihood below 0.75 were removed, gaps under 25 frames (about 1 s) were interpolated, and velocities were smoothed with a Savitzky–Golay filter [34].

From the tracked coordinates we defined five families of features describing behavioural microstructure: (i) investigation time, bout number and mean bout duration per stimulus (a bout is one continuous visit to an interaction zone), (ii) chamber occupancy and transitions, (iii) head orientation, defined as the angle between the ear-midpoint-to-nose vector and the vector to the stimulus centroid (facing: ≤45°; orientation index: its cosine), (iv) approach bouts (distance to the stimulus decreasing for ≥0.4 s by ≥2 cm), approach speed relative to baseline speed, and approach completion; and (v) total distance (for full definitions see Supplementary Methods S2, S3). To examine how behaviour unfolded over the session, features were analysed in 60-s bins across the 10-min test (see Supplementary Methods S4).

### 2.4. Behavioural decoding

To test whether these pose-derived features carry information about feeding state and housing history, we trained supervised classifiers (see Supplementary Methods S6). Random forests were trained on 60 pose-derived features from the 10-min session to classify feeding state and housing condition (n=49), with subgroup decoders within housing or feeding strata and a four-class decoder. Performance was estimated by 10× repeated stratified 5-fold cross-validation (one observation per mouse) and summarised as the area under the receiver operating characteristic curve (AUC) and balanced accuracy; significance was tested against 200 label permutations. Feature-family importance was the drop in balanced accuracy after jointly shuffling a family in held-out folds. For comparison, a decoder was trained on 16 manually scored measures with the same pipeline and partitions.

### 2.5. Immunofluorescence and activity-associated mapping

We used Zif268 immunofluorescence as a marker of neuronal activation in regions involved in social behaviour, feeding, and the integration of these processes, i.e., the PVN, SON, BNST, BLA, CeA. In the SON and PVN, assessment of neuronal activity was restricted to OXT neurons. Ninety minutes after testing, mice were anaesthetised with an intraperitoneal injection of ketamine (10%; 1 mL/kg; WDT, Garbsen, Germany) and xylazine (2%; 0.5 mL/kg; Serumwerk Bernburg AG, Bernburg, Germany). Subsequently, animals were perfused transcardially with PBS followed by 4% paraformaldehyde. Free-floating coronal sections (30 µm) were blocked in 5% normal goat serum and incubated with rabbit anti-EGR1/Zif268 (1:250; Cell Signaling, #4153) and mouse anti-oxytocin (OXT) (1:1,000; Dr Maurice Manning), followed by Alexa Fluor 594 goat anti-rabbit and Alexa Fluor 488 goat anti-mouse secondary antibodies (both 1:1,000; Invitrogen, #A11012, #A11001), and mounted with ROTI Mount FluorCare with DAPI. Images were acquired on a Leica DM6 B microscope. Zif268-positive nuclei and OXT-immunoreactive cells were counted semi-automatically in ImageJ v1.53c on 2–8 bilateral sections per region by a blinded investigator. Brain regions were defined by anatomical landmarks and the Allen Mouse Brain Atlas (mouse.brain-map.org) (see Supplementary Methods S7). Zif268 expression was quantified as the percentage of DAPI-positive cells expressing Zif268. Expression was compared across ST, FT and SFPT contexts within each group.

### 2.6. Statistical analysis

Analyses used GraphPad Prism (version 11.1.0; GraphPad Software, Boston, MA, USA) and Python. Normality was tested with the Shapiro–Wilk test and homogeneity of variances with Levene’s test. Housing × feeding effects were tested by two-way ANOVA and stimulus × treatment effects by two-way repeated-measures ANOVA, followed by Tukey’s multiple-comparisons test; sphericity was assessed and corrected where necessary. Planned comparisons were performed using two-sided unpaired Student’s *t*-tests. Preference indices were compared with zero by one-sample t-tests or Wilcoxon tests. Zif268 expression was compared across contexts within each group using one-way ANOVA followed by Tukey’s multiple-comparisons test (or Welch’s ANOVA followed by Games–Howell’s test, as appropriate). Data are mean ± SEM or individual points; significance was set at p<0.05. Figures were generated using GraphPad Prism, Python and BioRender and assembled using Affinity Designer (version 2.6.2; Serif Europe, Nottingham, UK). Statistical analyses relevant to pose-derived microstructure and temporal analysis here are described in the Supplementary Methods (S5).

## 3. Results

### 3.1. Feeding state and housing history each bias choice between social and food rewards

Habituation to the reward was comparable across groups (Fig. S1A–C). In the SFPT, stimulus-directed investigation was dependent on stimulus and treatment (F(3,45)=8.16, p<0.001; Fig. 1C). GH-AL and SH-AL mice displayed social preference behaviour and investigated the social stimulus more than the food stimulus (p=0.017 and p=0.002, respectively). In contrast, GH-FD mice preferred the food over the social stimulus (p=0.003), and SH-FD mice displayed no preference. In general, FD increased total investigation (not shown).

The preference index (Fig. 1D) was positive in GH-AL (t(9)=5.56, p<0.001) and SH-AL mice (t(10)=5.26, p<0.001) and negative in GH-FD mice (t(9)=−2.51, p=0.033), and not different from zero in SH-FD mice (t(17)=1.437, p=0.169). Two-way ANOVA revealed main effects of feeding state (F(1,45)=22.4, p<0.001) and housing (F(1,45)=7.36, p=0.009) without an interaction (F(1,45)=2.36, p=0.132), consistent with additive effects: FD shifted preference toward food, and single housing shifted it toward the social stimulus. Planned comparisons localised these effects. FD shifted preference toward food in both GH mice (GH-AL vs. GH-FD: t(18)=4.84, p<0.001) and SH mice (SH-AL vs. SH-FD: t(27)=2.26, p=0.032), and the food bias was stronger in GH-FD than in SH-FD mice (t(26)=2.74, p=0.011).

Control tests provided evidence against general group differences in naturally occurring preferences: all groups preferred a social stimulus over an empty cage, and food over an empty cage (except SH-AL mice). Also, results from the CPP test confirmed that Honey Pops are rewarding (Fig. S2). As the total distance travelled was found to be lower in FD mice (F(1,45)=30.3, p<0.001 vs. AL) and higher in SH mice (F(1,45)=9.77, p=0.003 vs. GH; Fig. 1I), locomotion has been accounted for in the kinematic analyses below. Overnight food deprivation reduced body weight (Fig. 1J), and weight loss did not correlate with preference in any group (not shown).

### 3.2. Hunger and isolation reshape engagement through distinct behavioural routes

A food bias can arise from more frequent food-chamber entries or from longer engagement after entry. Neutral-to-stimulus transitions did not differ between groups (Fig. 1G), and GH-FD mice made no more food-chamber entries despite investigating food more. Instead, food-directed bouts were longer in GH-FD than in GH-AL mice (Stimulus × Treatment, F(3,45)=3.02, p=0.039; p=0.006; Fig. 1E), and social bouts were fewer after food deprivation (GH, p<0.001; SH, p=0.040; Fig. 1F). Time in the chamber added a related but distinct measure (Stimulus × Treatment, F(3,45)=12.9, p<0.001; Fig. 1H): SH-FD mice spent more time in the social than in the food chamber (p=0.033), despite showing no difference in investigation.

Approach kinematics separated the two manipulations (Fig. 2A–C). Social-approach speed showed a time × treatment interaction (F(3,45)=8.68, p<0.001; Fig. 2B), rising above baseline in SH-FD mice (p<0.001). Food-approach also showed a time × treatment interaction (F(3,45)=6.83, p<0.001; Fig. 2C), rising above baseline in both GH-FD and SH-FD mice (both p<0.001) and was highest in SH-FD mice (p<0.001 versus GH-FD). SH-FD mice therefore approached both stimuli faster than any other group, in contrast to the food-directed shift in time spent near the stimuli observed in GH-FD mice.

**Figure 2.**
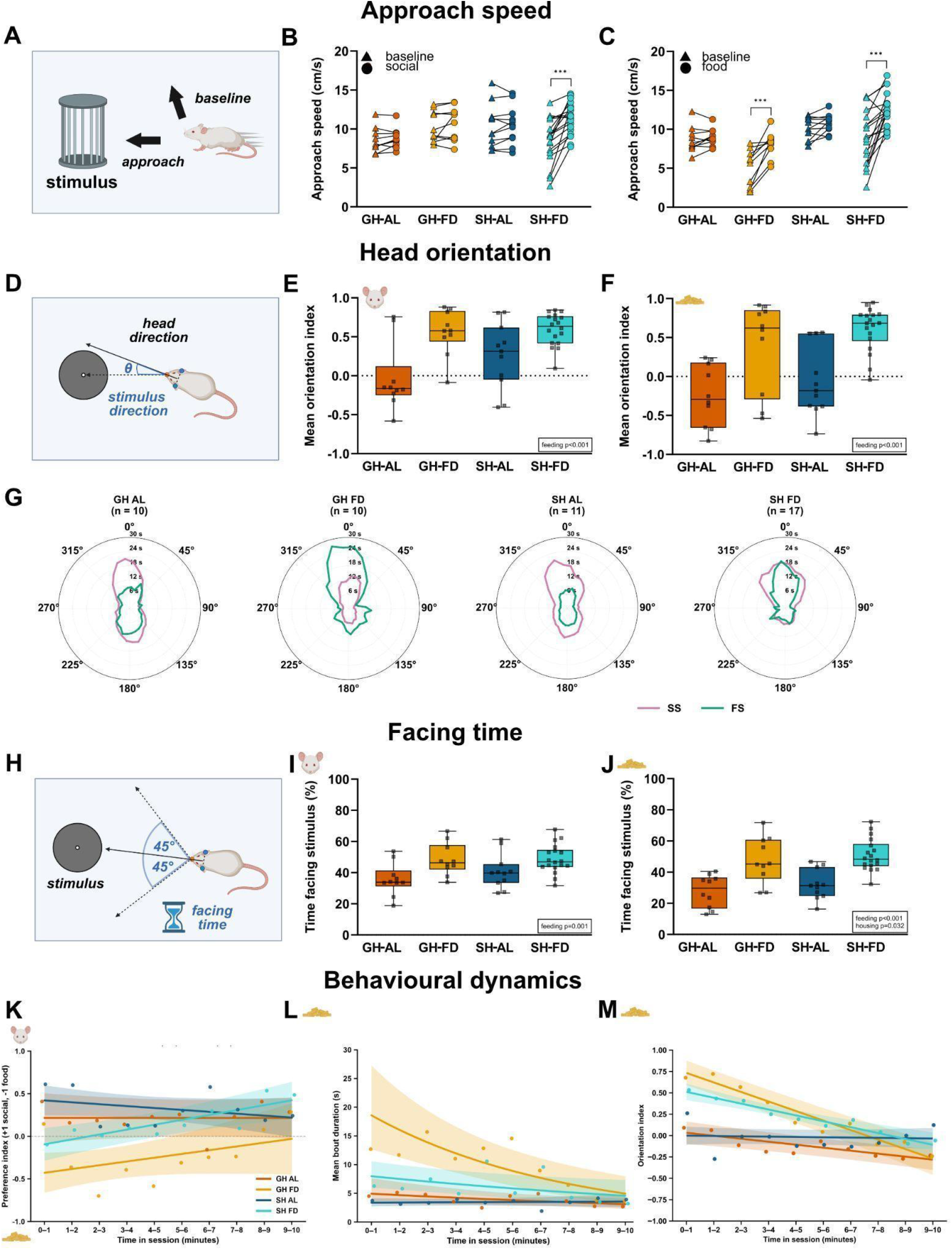
Hunger and isolation reshape engagement through distinct behavioural routes. (A) Schematic of stimulus-directed approach speed. (B,C) Approach speed toward the social (B) and food (C) stimulus relative to baseline in GH-AL, GH-FD, SH-AL and SH-FD mice. (D) Schematic of head orientation. (E,F) Orientation index toward the social (E) and food (F) stimulus. (G) Polar plots of head orientation. (H) Schematic of facing (within 45° of the stimulus). (I,J) Proportion of time facing the social (I) and food (J) stimulus. Data are presented as individual data points or box-and-whisker plots (minimum to maximum, all points). Statistical analysis: ***p<0.001 (B) and (C) (Two-way repeated-measures ANOVA, Tukey post hoc tests); (E), (F), (I), and (J) (Two-way repeated-measures ANOVA). (K) Model-fitted preference index over the session, by treatment group (GEE fit; ribbons95% CI; dotsraw bin means). (L) Model-fitted mean food investigative bouts over the session, by treatment group (GEE fit; ribbons95% CI; dotsraw bin means). (M) Model-fitted orientation index over the session, by treatment group (GEE fit; ribbons95% CI; dotsraw bin means). Sample sizes: GH-AL, n=10; GH-FD, n=10; SH-AL, n=11; SH-FD, n=18. Abbreviations: AL, fed ad libitum; FD, food-deprived; GH, group-housed; SH, single-housed. (A,D,H) Created in BioRender.

Feeding state altered head orientation toward the social (F(1,45)=23.1, p<0.001; Fig. 2E) and food (F(1,45)=29.2, p<0.001; Fig. 2F) stimulus and facing time toward each (social: F(1,45)=12.1, p=0.001; food: F(1,45)=34.5, p<0.001; Fig. 2I,J), with an additional housing effect on facing time toward food (p=0.032). Food-approach completion depended on feeding state (F(1,45)=9.11, p=0.004) and housing (F(1,45)=4.55, p=0.038), and completed food approaches lasted longer in GH-FD than GH-AL and SH-FD mice (both p<0.001; Feeding × Housing interaction F(1,45)=5.07, p=0.029). Thus, FD increased food-directed orientation and sustained engagement, whereas SH additionally increased approach speed toward both stimuli. A null preference score in SH-FD mice therefore coexisted with active, rapid engagement rather than disengagement.

Both FD (p<0.001) and GH (p=0.001) shifted preferences away from the social zone, but only GH-FD animals showed a net preference for food (mean index −0.26). This food preference declined across the session, with hungry animals increasing their relative social preference over time (+0.062 per min, p=0.004) (Fig. 2K). Correspondingly, FD lengthened visits to the food zone (p<0.001), elevating geometric mean bout durations to a peak of 8.3 s in GH-FD mice compared to 3.2-3.4 s in AL mice. Group housing amplified bout duration exclusively during hunger, resulting in 1.5-times longer visits in GH-FD than SH-FD animals (p=0.014)—a compounding effect driven entirely by bouts in the early half of the session (p=0.028) before disappearing later on in the session (p=0.88) (Fig. 2L).

Orientation toward the food stimulus changed dynamically across groups over the course of the session (Treatment × Time interaction: χ^2^ (3)=88.5, p<0.001; global Wald p<0.001). Food orientation decreased over time in GH-AL, GH-FD, and SH-FD mice, with the steepest decline observed in group-housed food-deprived (GH-FD) animals. Relative to GH-AL mice, this downward trajectory was significantly steeper in both GH-FD (β = −0.077, *p_Holm_* = 3. 5×10^−6^) and SH-FD mice (β= −0.034, *p_Holm_* = 0.045). In contrast, SH-AL mice exhibited a significantly less pronounced decline over time (β= +0.031, *p_Holm_* = 0.030) (Fig. 2M).

### 3.3. Machine-learning decoding of pose captures state-specific behavioural signatures

Cross-validated decoders performed above chance (Fig. 3 A–D).

**Figure 3.**
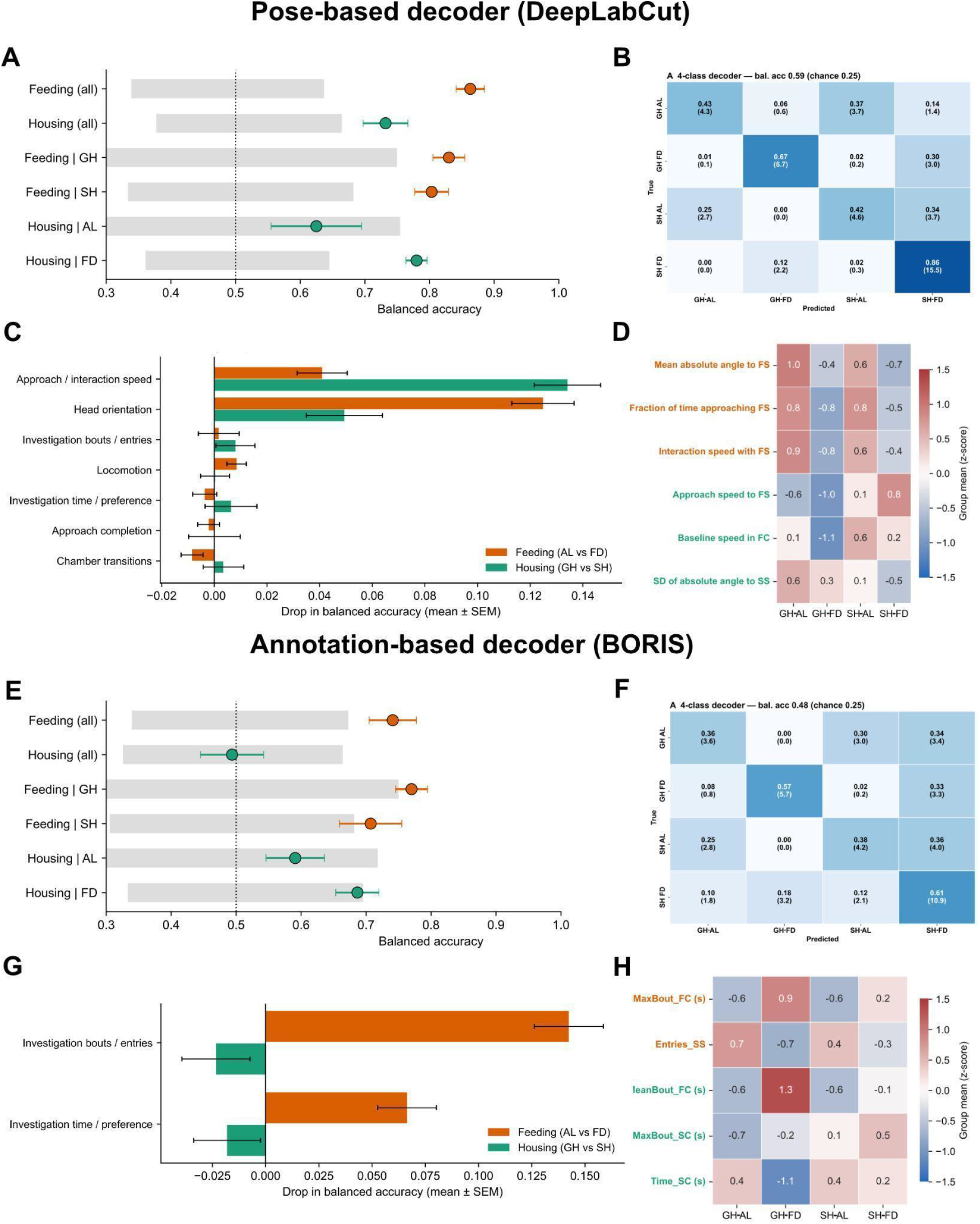
Machine-learning decoding of pose captures state-specific behavioural signatures. (A) Cross-validated balanced accuracy across decoding tasks. (B) Four-class confusion matrix. (C) Feature family importance for feeding and housing decoders. (D) Group mean z-scores for top-ranked features. (E) Cross-validated balanced accuracy for manual metrics. (F) Four-class confusion matrix for manual metrics. (G) Metric family importance for manual decoders. (H) Group mean z-scores for top manual metrics.

Feeding state was decoded with a higher AUC (AUC=0.94, p≤0.005, n=49) than housing condition (AUC=0.85, p≤0.005). Feeding-state decoding held within GH (AUC=0.94, n=20) and SH mice (AUC=0.91, n=29), whereas housing was decoded under food deprivation (AUC=0.86, p=0.010, n=28) but not significantly under AL conditions (AUC=0.70, p=0.114, n=21). Balanced accuracy was 0.86 for feeding state and 0.73 for housing (Fig. 3A).

Different feature families carried each state (Fig. 3C). The feeding decoder relied mainly on head orientation (accuracy drop 0.125 ± 0.012) and secondarily on approach and interaction speed (0.041 ± 0.009). The housing decoder relied mainly on approach and interaction speed (0.134 ± 0.012) followed by head orientation (0.049 ± 0.015). Group profiles (Fig. 3D) showed larger angles to food in AL mice and faster food approach and higher baseline speed in SH mice. A four-class decoder identified all groups (balanced accuracy 0.59, chance 0.25; AUC=0.88), with errors almost exclusively between housing conditions within the same feeding state.

A decoder trained only on manually scored measures (BORIS) predicted feeding state (AUC=0.84, p≤0.005) but did not decode above chance housing (AUC=0.56, p=0.463; Fig. 3E–H). Thus, manually scored event counts and durations did not capture the housing-related differences that spatial and kinematic features resolved.

### 3.4. Neuronal recruitment during social versus food choice depends on internal state and social history

We compared the Zif268 expression during the SFPT with that after the social-only (ST) and food-only (FT) tests within each group (Fig. 4). This comparison identifies regions that are differentially recruited, when social and food stimuli are available simultaneously, relative to either stimulus alone.

**Figure 4.**
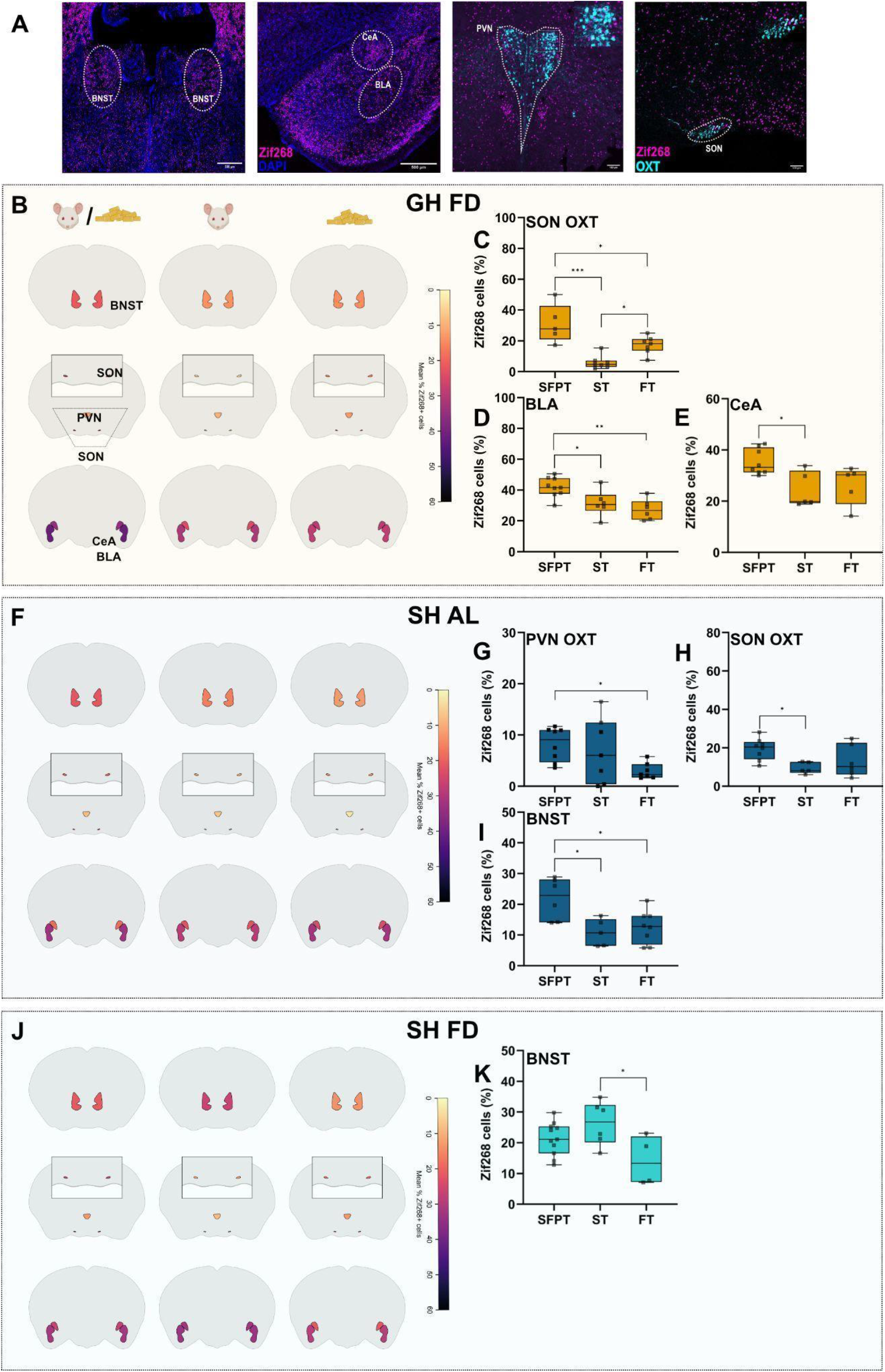
Neuronal activity patterns in response to social versus food choices depend on internal state and housing conditions. (A) Representative DAPI, OXT and Zif268 images for BNST, BLA, CeA, PVN-OXT and SON-OXT. (B-E) GH-FD mice: sections across social-only, food-only and SFPT contexts, and quantification in SON-OXT (C), BLA (D) and CeA (E). (F-I) SH-AL mice: sections across social-only, food-only and SFPT contexts, and quantification in PVN-OXT (G), SON-OXT (H), and BNST (I). (J-K) SH-FD mice: sections across social-only, food-only and SFPT contexts, and quantification in BNST (K). Data are presented as box-and-whisker plots (minimum to maximum, all points). *p<0.05, **p<0.01, ***p<0.001 (One-way ANOVA, Tukey’s test). Sample sizes: n=5-11. Abbreviations: AL, fed ad libitum; BLA, basolateral amygdala; BNST, bed nucleus of the stria terminalis; CeA, central amygdala; DAPI, 4′,6-diamidino-2-phenylindole; FD, food-deprived; GH, group-housed; OXT, oxytocin; PVN, paraventricular nucleus; SFPT, social versus food preference test; SH, single-housed; SON, supraoptic nucleus;; Zif268 (EGR1)=early growth response 1.

In GH-AL mice, no region differed between contexts (data not shown). In GH-FD mice (Fig. 4B), context affected Zif268 expression only in SON OXT neurons (F(2,16)=15.6, p<0.001), which was higher after exposure to the SFPT than the ST (p<0.001) or FT (p=0.019). Neuronal activity in the BLA was also higher in response to the SFPT than the ST (p=0.031) or FT (p=0.003; F(2,18)=8.55), whereas in the CeA Zif268 expression was higher after exposure to the SFPT than the ST (p=0.022; F(2,15)=5.58).

In SH-AL mice (Fig. 4C), PVN OXT neuronal activity differed across contexts (F(2,10.5)=8.20, p=0.007) and was higher during the SFPT than during the FT (p=0.007). Similarly, SON OXT activity differed across contexts (F(2,16)=4.27, p=0.033) and was higher in the SFPT than in the ST (p=0.032). BNST expression also differed (F(2,16)=6.51, p=0.009) and was higher in the SFPT than in both ST (p=0.014) and FT (p=0.020). In SH-FD mice (Fig. 4D), only BNST expression differed across contexts (F(2,18)=4.33, p=0.029), being higher in the ST than in the FT (p=0.029).

## 4. Discussion

Hunger is known to shift social–food preference [10,12–14, 35]. Here, we show that machine-learning-based, pose-derived microstructure resolves the components of social–food choice that a single preference score, or manual scoring, merge. This shift is assembled from separable components that carry information about internal state beyond a simple preference score. Four findings stand out. First, FD increased the duration mice are engaged with the food stimulus, but did not alter the choice of the chamber entered. Second, SH attenuated the FD-induced food shift: although SH-FD mice did not display any preference (near-zero preference index), they approached both stimuli faster than any other group. Third, pose-derived features decoded housing history, which manually scored measures did not. Fourth, simultaneous presentation of both rewards was associated with state-specific Zif268 patterns in amygdalar and hypothalamic regions, with increased neuronal activity in the SON, BLA and CeA in GH-FD mice and in the PVN, SON and BNST in SH-AL mice. Together, these findings indicate that behavioural microstructure adds interpretable information beyond a single preference parameter.

### 4.1. Preference is expressed in dwell and engagement, not in the choice of destination

A preference score (Fig. 1D) can arise in two ways: animals can choose one stimulus more often, and/or for longer. Our data point to the second. Transitions from the neutral chamber to either stimulus chamber did not differ between groups, and the probability of repeated visits to the same chamber was similar across feeding and housing conditions (Fig. 1G). In other words, hungry and fed mice, independent of housing conditions, were equally likely to approach and investigate the food or social chamber, and to return to the one they had just left. However, they differed in the duration of investigating each stimulus with GH-FD mice showing lengthened food-investigation bouts (Fig. 2L), reduced the number of social bouts (Fig. 1F), and increased food-directed orientation and facing time (Fig. 2F,J), while the number of entries into the food chamber did not increase (Fig. 1G) compared with GH-AL mice. The preference shift, therefore, reflects a reallocation of time and attention within visits, not a change in the stimulus choice.

Among AL-fed mice, GH and SH animals did not clearly differ in social or food investigation, chamber time or preference index (Fig.1, 2), indicating that housing alone did not shift behaviour in the SFPT. Housing mattered when combined with FD: SH-FD mice approached both stimuli faster than any other group (Fig. 2B,C) and spent more time in the social than in the food chamber (Fig. 1H), despite no difference in stimulus-directed investigation (Fig. 1C). Thus, chamber occupancy, investigation time, bout structure and approach speed each capture a different aspect of behaviour. They are not interchangeable measures of preference. A non-significant preference score also does not show that the two rewards were valued equally, so we do not interpret the SH-FD result that way. These measures describe how mice approach and engage with the stimuli, not how motivated they subjectively are.

### 4.2. A need-based landscape as a framework for the time course

The temporal profile of food-directed behaviour suggests a way to organise our findings within the need-based energy landscape introduced above, which may explain why dwell and engagement, but not the choice of destination, differed between groups. In FD mice, food-investigation bouts and the orientation index toward food were highest early in the session and declined over time, converging towards the values of AL mice by the end (Fig. 2L,M). Social-directed engagement remained comparatively stable (not shown). This pattern resembles the organisation of choice reported in mice facing conflicting hunger and thirst, in which choices form persistent bouts with stochastic switches between wells [15].

Our observations are compatible with this view in broad terms (Fig. 5). If hunger initially deepens the food well, a GH-FD mouse that reaches the food would tend to stay engaged with it, lengthening food bouts and holding orientation toward food, which is the pattern seen early in the session. As food is consumed and the need is addressed, the well flattens and food-directed engagement would converge toward that of fed mice. Social engagement shaped by a lower although steadier need in GH mice, would change little over the same period. On this reading, hunger is the more urgent need and is addressed first, whereas social engagement is maintained at a lower level. In this framework, need shapes how long a state is held once it is adopted, which fits our finding that destination probabilities were similar across groups while dwell and engagement differed. For SH-FD mice, one possibility is that isolation deepens the social well while hunger deepens the food well, so that two strong pulls produce rapid approach to both stimuli and a near-balanced index. We offer this as a hypothesis.

**Figure 5.**
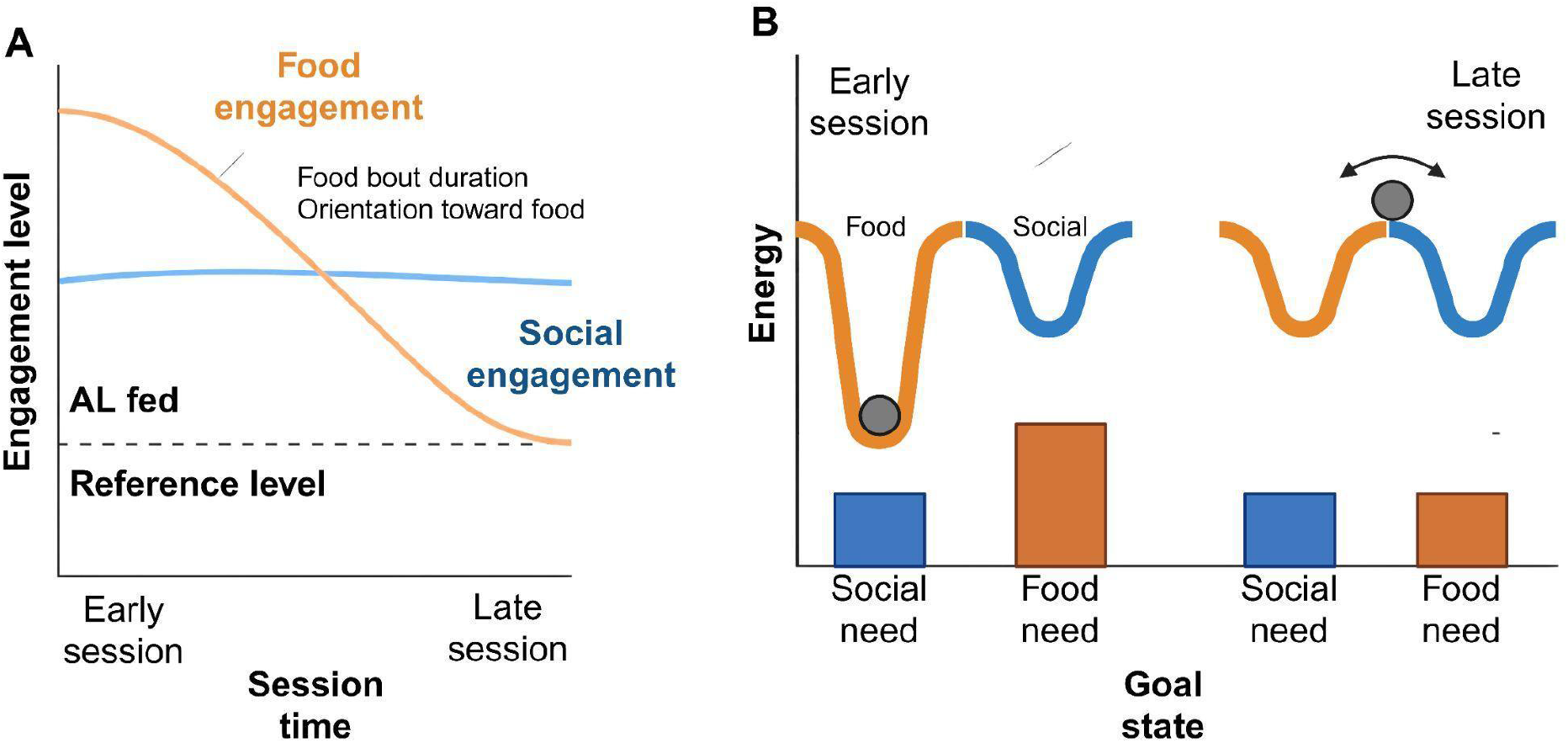
Conceptual framework for the time course of food-directed engagement. (A) Predicted time course of engagement. In food-deprived mice, food-directed engagement (food-investigation bout duration and orientation toward food) is highest early in the session and declines, converging towards the reference level of ad libitum (AL) fed mice. Social engagement is assumed to remain comparatively stable. (B) Need-based energy landscape. Each goal state (food, social) is represented as a well, and the ball represents the animal’s current goal state. The depth of a well reflects the strength of the corresponding need (bars). Early in the session, the high food need creates a deep food well that holds the animal in persistent food engagement. As the food need is satisfied, the well flattens, the food and social needs become comparable, and the animal can switch between goal states, so engagement converges towards that of fed mice. This framework is offered as a hypothesis and was not fitted or tested here. Adapted from Richman et al. (2023).

We present this as an interpretive framework, not a tested model. Unlike the head-fixed mice with equidistant rewards and hundreds of choices per session in the original hunger-thirst study [15], our mice move freely between distant stimuli and contribute few excursions in 10 min, so we could not fit or test a landscape model. We also measured investigation, which includes sniffing and eating, and not consumption directly, so we cannot say that the decline in food engagement reflects satiation. Food orientation also fell in ad libitum mice, so habituation to the stimuli may contribute, and the stability of social engagement over 10 min does not show that social need cannot be satiated.

### 4.3. Pose-derived behaviour can distinguish state-related signatures

The decoding analyses show that high-dimensional behaviour contains information that a preference index lacks (Fig. 3). Feeding state was classified with high accuracy, with head orientation being the most influential behaviour. Housing condition was more closely tied to approach and interaction speed, especially under FD but not AL feeding conditions, which suggests that hunger may unmask housing-related differences in approach. However, a significant and a non-significant decoder do not constitute a test of interaction, and the subgroups differ in size (AL, n=21; FD, n=28). The two manipulations nonetheless left partly distinct signatures within one assay, in line with the univariate results: feeding state altered orientation and facing time, whereas single housing altered approach speed in SH-FD mice.

Pose-based analysis captured more of the information in behaviour than manual scoring did. Both approaches decoded feeding state, but pose-derived features did so more accurately (AUC 0.94 vs 0.84), and only pose-derived features decoded housing history (AUC 0.85 vs 0.56). Continuous spatial and kinematic measures, such as head orientation and approach speed, register differences in how animals engage with a stimulus that event counts and bout durations miss. This is a practical advantage: housing-related differences that manual scoring would have missed were recoverable from the same videos. Feature importance indicates which inputs support a model within a given pipeline and does not identify mechanistic drivers, and with 49 animals the performance estimates remain uncertain, so generalisation to held-out cohorts, other strains and other rewards still needs to be tested.

### 4.4. Interpreting the neuronal activity-associated findings

Simultaneous availability of social and food stimuli was associated with Zif268 patterns that differed from those after either stimulus alone, and the regions involved were dependent on feeding state and housing (Fig. 4). In GH-FD mice, choice was accompanied by higher activity in the BLA, CeA, and SON OXT neurons. The BLA contains separable representations of social and sucrose reward [21] and neurons that control both feeding and social interaction [22], which makes it a plausible site for resolving competition between the two stimuli. In SH-AL mice, expression was higher in the BNST and in PVN and SON OXT neurons. Notably, this housing-related difference in neuronal recruitment was not accompanied by a clear behavioural difference between SH-AL and GH-AL mice, indicating that housing history can change how the brain engages during choice without changing the overt behavioural output measured here. OXT signalling has been linked to the isolation-dependent reorganisation of social–food priorities in rats [14], which makes the OXT findings in SH-AL mice a candidate mechanism for this dissociation. In SH-FD mice, no choice-specific increase was detected. The absence of a significant difference does not establish the absence of recruitment, and the heterogeneity across groups is not consistent with a single, uniformly activated circuit for social–food prioritisation [35].

### 4.5. Limitations

Our conclusions apply to adult male outbred CD-1 mice, one palatable food reward and a 16-h deprivation protocol; whether the same strategies appear in females, other strains and other rewards remains to be tested. Single housing combines reduced social contact with other consequences of isolation, so we interpret it as a housing history and not as a selective manipulation of social motivation. The SH-FD group was larger than the others, and group-wise inferences were made with methods that account for unequal variances. Approach speed was normalised to each animal’s baseline speed, and the main effects of feeding state and housing on distance were modelled explicitly, which limits but cannot exclude a locomotor contribution to the kinematic results. Food investigation includes sniffing and eating, and no direct food intake measure was taken, so the temporal decline in food engagement cannot be attributed to satiation, and the landscape framework in Section 4.2 is offered as an interpretation. The decoders were validated by repeated cross-validation with label permutation in 49 animals; because the manually scored decoder used fewer measures and the contrast between decoders was not tested directly, the comparison is a benchmark of information content, and confirmation in independent cohorts is the next step. Zif268 is an indirect, time-averaged marker of activation, and the SFPT differs from single-stimulus tests in the number of stimuli as well as in choice, so these data identify regions associated with dual-stimulus availability and do not establish a causal role in prioritisation. Our design identifies differential recruitment during choice, meaning regions that respond differently when social and food stimuli are available together than when either is presented alone. It does not measure absolute activation relative to an unstimulated baseline. Regions engaged similarly across all three contexts would not appear as differentially recruited, so a null result does not exclude their involvement in choice-related processing.

## Conclusion

Pose-derived microstructure turns a single preference score into a description of the behavioural strategy behind it: how animals approach, orient towards and stay engaged with each option. It shows that feeding state and housing history act on different components of this strategy, with feeding state mainly altering orientation and engagement and single housing altering approach speed. It also shows that markerless pose estimation captures information about internal state that conventional manual scoring does not. Reporting such measures alongside preference scores would make established assays more informative and would help identify the processes, and ultimately the circuits, that choice assays are designed to probe.

## Declarations

The authors have no conflicts of interest to declare.

## Funding

This study was supported by the German Research Foundation (DFG) through the Neuroscience Graduate Programme “Neurobiology of Emotion Dysfunctions” (GRK 2174, to V.R. and I.N.).

## Acknowledgements

We thank Dr. Rohit Menon and Dr. Oliver Bosch for scientific discussion, and M. Fuchs, A. Havasi and R. Maloumby for excellent technical help.

## CRediT authorship

AS: Formal analysis, Visualization, Writing – Original Draft; NB: Investigation, Data Curation, Visualization; SP: Formal analysis, Visualization, Writing – Original Draft; IDN: Resources, Supervision, Project administration, Funding acquisition, Writing – Review & Editing; VR: Conceptualization, Methodology, Investigation, Validation, Data Curation, Visualization, Supervision, Project administration, Funding acquisition, Writing – Original Draft, Writing – Review & Editing.

## Data availability

The analysis code, including the scripts for pose-derived feature extraction, decoding and statistical analysis, is available at https://github.com/aryanatulsurana-hue/pose-features-decoder and archived on Zenodo (DOI: <u>10.5281/zenodo.24138305</u>) under the MIT licence. The source data underlying the figures and the trained DeepLabCut model, per-frame tracking files are available from the corresponding author upon reasonable request.

## Generative AI declaration

During manuscript preparation, AI (ChatGPT, Claude) was used solely for language editing. The authors reviewed, edited and take full responsibility for the content.

## Notes

https://github.com/aryanatulsurana-hue/pose-features-decoder

https://zenodo.org/records/24138306

### Competing Interest Statement

The authors have declared no competing interest.

