## Supplementary Information for "Pose-derived microstructure resolves how hunger and social isolation shape social–food choice in male mice"

### S1. Habituation and conditioned place preference

**Habituation.** Mice completed eight 5-min habituation sessions across three consecutive days. A Honey Pops pellet was placed in one end chamber, with its location counterbalanced across sessions. Latency to approach and consume the reward and total eating time were scored manually.

**Conditioned place preference (CPP).** Reward value was assessed in a two-chamber apparatus (54 × 54 × 41 cm) with distinct visual cues (vertical lines versus crosses) and tactile cues (LEGO® base plates). After a 15-min pretest, mice received six days of conditioning with two 30-min sessions per day, pairing Honey Pops and standard food with different chambers. Chamber assignment was counterbalanced for baseline preference. During the 15-min test, no food was present. The CPP index was [time in the Honey Pops-paired chamber − time in the standard-food-paired chamber] / [time in both chambers].

### S2. Chamber and stimulus zones

Custom Python scripts defined chamber boundaries and stimulus-interaction regions. Interaction regions were delineated manually around each stimulus cage and expanded by a factor of 1.85. Chamber occupancy required both the head and nose coordinates to lie within the chamber boundary. Stimulus investigation was scored when the nose entered the corresponding interaction region. The neutral chamber was the central chamber connecting the two end chambers.

### S3. Definitions of pose-derived measures

**Investigation.** An investigation bout was one continuous visit to a stimulus-interaction zone. Mean bout duration and bout count were computed separately for social and food investigation. The preference index was (social investigation − food investigation) / (social investigation + food investigation).

**Head orientation.** Head direction was the vector from the midpoint between the ears to the nose. Orientation was the angle between this vector and the vector from the nose to the centroid of the stimulus polygon. Absolute angles of 45° or less were classed as stimulus-facing, and facing time was the proportion of time spent facing. The orientation index was the cosine of the heading angle, from +1 (directly towards the stimulus) through 0 (perpendicular) to −1 (directly away).

**Approach.** An approach bout was detected when the signed distance to the stimulus edge decreased for at least 0.4 s and the mouse approached by at least 2.0 cm within the relevant chamber. A bout ended at the local distance minimum or when the mouse left the chamber. Approach-initiation rate was expressed per minute spent in the chamber. Approach-completion rate was the proportion of initiated approaches that reached the interaction zone.

**Speed.** A 6-cm radius around each stimulus defined an interaction-proximity zone, and frames in this zone were excluded from approach- and baseline-speed classification. Approach speed was calculated during stimulus-oriented movement towards the stimulus (radial speed above 0.5 cm/s), outside the proximity zone and with body speed above 1.0 cm/s. Baseline speed was calculated from all other in-chamber frames outside the proximity zone. Total distance travelled was computed from the body-centre trajectory.

**Chamber transitions and choice history.** Choice points were visits to the neutral chamber between successive end-chamber visits, and the next end chamber entered defined the choice. The number of choice points and the proportion directed to the food chamber were quantified. Transition probabilities between chambers were also computed. Choice history was the phi coefficient between the previous and next chamber:

$$\varphi = (ad - bc) / \sqrt{[(a + b)(c + d)(a + c)(b + d)]}$$

where a, b, c and d are the numbers of choice points going from social to social, social to food, food to social and food to food, respectively. Positive values indicate returning to the previously visited chamber and negative values indicate switching.

**S4. Temporal analyses**

Sessions were divided into 60-s bins. The preference index was calculated in each bin from time in the social and food zones. Bout duration was assigned to the bin in which the bout began, and bins without bout onsets were excluded from bout-duration analyses. To compare early and late food-investigation bouts, each animal's bouts were split at its median bout-onset time, giving equal numbers of bouts in each half.

**S5. Statistical models**

**Per-animal analyses.** Per-animal values were analysed with Housing × Feeding state ANOVAs (Type II sums of squares). Bout durations were log-transformed and choice counts transformed as log(1 + x). Group means of bout duration were geometric means of per-animal means, and preference indices were summarised as arithmetic means. Bout-duration effects were expressed as ratios of geometric means with 95% confidence intervals. Early–late comparisons used the same approach. Robustness was assessed with heteroscedasticity-consistent HC3 F-tests and Freedman–Lane permutation tests. Interaction and main-effect p values were unadjusted.

**Generalised estimating equations (GEEs).** Temporal changes in preference index, food-bout duration and orientation index were analysed with GEEs, with animals as clusters and an exchangeable working correlation. Preference and orientation indices used a Gaussian distribution with identity link, and bout durations used a Gamma distribution with log link. Each model included housing, feeding state, time and all interactions. Housing (GH=+0.5, SH=−0.5) and feeding state (FD=+0.5, AL=−0.5) were effect-coded and time was centred at mid-session, so main effects are differences at mid-session averaged over the other factors and the time effect is the average slope across groups. Standard errors used a small-sample, bias-reduced sandwich estimator and coefficients were tested with t-distributions. Group-specific slopes and pairwise slope contrasts were derived from the models, with Holm correction across the four slopes or four contrasts per outcome. Fitted trajectories were linear on the link scale. Bout-duration analyses were repeated at the individual-bout level. Robustness was checked with alternative correlation structures and random-slope mixed models.

**Stimulus-choice (excursion) model.** Each bout sequence was divided into excursions, spanning departure from one stimulus zone to entry into the next, including direct re-entries into the same zone. The binary outcome was whether the next zone entered was the social zone. Relative session progression was (rank − 0.5) / n, where n is the number of excursions. A binomial GEE with logit link, animals as clusters, an exchangeable working correlation and bias-reduced standard errors modelled social-zone entry as a function of previous zone, housing, feeding state, relative time and all interactions. Predictors were sum-coded and time was centred at its mean. Effects involving the previous zone quantified return tendency, the log odds ratio of entering the social zone after leaving the social rather than the food zone (positive: returning to the zone just left; negative: switching). Model terms were tested with unadjusted Wald $\chi^2$ tests. For each group, predicted probabilities of social-zone entry were averaged across previous zones, weighted by the observed proportion of excursions originating from the social zone. Probabilities at the first and last excursion and their difference were estimated with delta-method 95% confidence intervals and Holm correction across groups.

### S6. Behavioural decoding

**Tasks.** Binary random-forest decoders classified feeding state (AL versus FD) and housing condition (GH versus SH) across all mice (n=49). Subgroup analyses decoded feeding state within housing conditions (GH, n=20; SH, n=29) and housing within feeding states (AL, n=21; FD, n=28). A four-class decoder classified the four housing-by-feeding groups and was scored with the macro-averaged one-vs-rest AUC.

**Features.** Decoders used 60 pose-derived features from the 10-min SFPT session, in the families head orientation, approach and interaction speed, investigation bouts and entries, locomotion, investigation time, approach completion and chamber transitions. Features that were constant, empty or highly similar to one another were removed before modelling, for eg: Orientation Index and Angle to stimulus.

**Cross-validation.** Performance was estimated by 10× repeated stratified 5-fold cross-validation. Each mouse contributed one observation, so no animal appeared in both training and test folds. AUC was computed from out-of-fold class probabilities and averaged over repeats, and balanced accuracy was also cross-validated.

**Permutation tests.** Balanced accuracy was compared with label-permutation null distributions generated by shuffling labels 200 times and repeating the full cross-validation pipeline for each shuffle. P values were (1 + number of null values ≥ observed) / (1 + 200), so the smallest attainable value is about 0.005.

**Feature-family importance.** Importance of each feature family was the decrease in balanced accuracy on held-out folds after jointly shuffling all columns of that family across test animals (5 shuffles per fold, averaged across folds). Z-scored group means described the direction of key features in the four groups.

**Manual-scoring comparison.** A separate decoder was trained on 16 manually scored (BORIS) measures with the same pipeline, hyperparameters and cross-validation settings (random seed 0), so both decoders used identical animal-level partitions and neither was tuned.

**S7. Immunofluorescence quantification**

**Regions of interest.** ROIs were defined with anatomical landmarks and the Allen Mouse Brain Atlas (AMBA; mouse.brain-map.org) and had consistent dimensions across animals and sections.

| Region | AMBA levels | ROI definition |
|---|---|---|
| BNST | 54–57 | 750 × 1,300 µm oval, positioned relative to the anterior commissure and internal capsule |
| BLA | 66–71 | 340 × 600 µm oval, located using the optic tract, external and amygdalar capsules and substantia innominata |
| CeA | 66–71 | 700 × 800 µm oval, same landmarks as BLA |

| Region | AMBA levels | ROI definition |
| --- | --- | --- |
| PVN | 58–63 | Manually outlined using anatomical landmarks and OXT immunofluorescence |
| SON | 58–63 | Identified relative to the optic tract and OXT immunofluorescence |

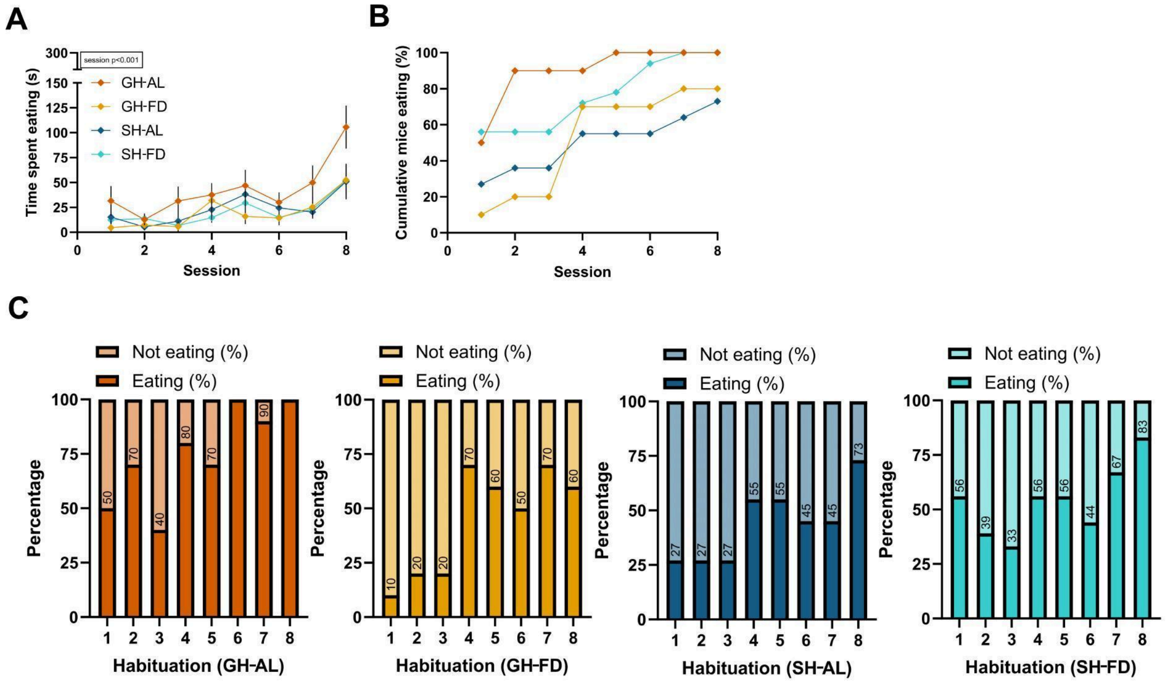


**Supplementary Figure 1. Food-reward habituation.** (A) Time eating the food reward across eight habituation sessions in GH-AL, GH-FD, SH-AL and SH-FD mice (Two-way repeated-measures ANOVA, main effect of Session: $F(3.81, 175)=17.7$, $p<0.001$, Greenhouse–Geisser $\varepsilon=0.544$). (B) Cumulative percentage of mice that ate the food reward at least once (C) Percentage of mice eating per session per group; numbers indicate percentages. Sample sizes: GH-AL, n=10; GH-FD, n=10; SH-AL, n=11; SH-FD, n=18. Abbreviations: AL, fed ad libitum; FD, food-deprived; GH, group-housed; SH, single-housed.

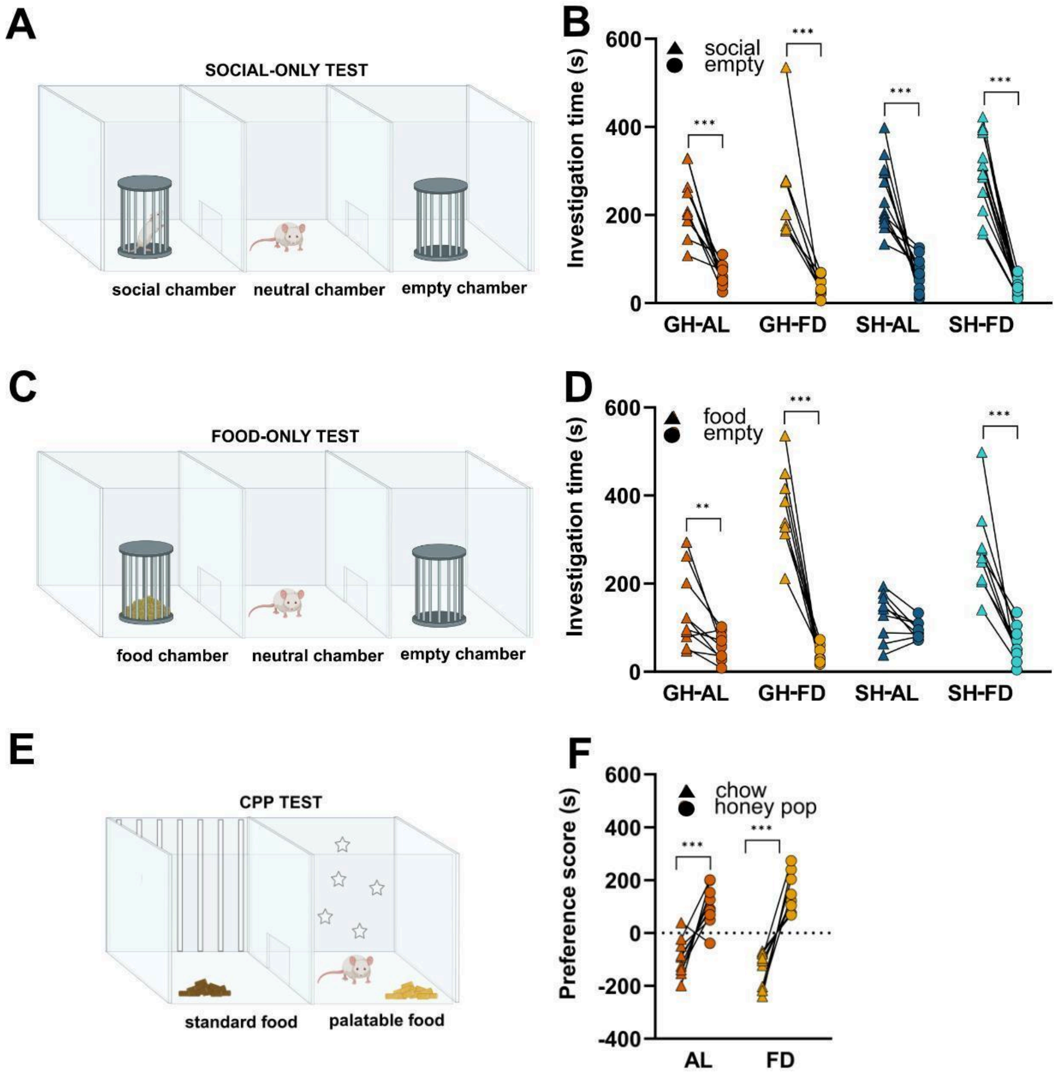


**Supplementary Figure 2. Control behavioural tests.** (A) Social-only test design. (B) Investigation time toward social vs. empty cage in GH-AL, GH-FD, SH-AL and SH-FD mice (main effect of Stimulus: $F(1,39)=167$, $p<0.001$, Two-way repeated-measures ANOVA). (C) Food-only test design. (D) Investigation time toward Honey Pops vs. empty cage (Stimulus × Treatment interaction: $F(3, 32)=19.0$, $p< 0.001$; Two-way repeated-measures ANOVA; **$p<0.01$, ***$p<0.001$, Tukey post hoc). (E) CPP test design. (F) CPP preference index: [time in Honey Pops-paired chamber − time in standard food-paired chamber] / total time in both chambers (main effect of chamber $F(1,18)=72.5$, $p<0.001$, Two-way repeated-measures ANOVA). Data are presented as individual data points. Sample sizes: n=8-13. Abbreviations: AL, fed ad libitum; FD, food-deprived; GH, group-housed; SH, single-housed.
